# Mosquito midgut stem cell cellular defense response limits *Plasmodium* parasite infection

**DOI:** 10.1101/2023.08.02.551669

**Authors:** Ana-Beatriz F. Barletta, Jamie C. Smith, Emily Burkart, Simon Bondarenko, Igor Sharakhov, Frank Criscione, David O’Brochta, Carolina Barillas-Mury

## Abstract

A novel cellular response of midgut progenitors (stem cells and enteroblasts) to *Plasmodium berghei* infection was investigated in *Anopheles stephensi*. The presence of developing oocysts triggers proliferation of midgut progenitors that is modulated by the Jak/STAT pathway, and proportional to the number of oocysts on individual midguts. The percentage of parasites in direct contact with enteroblasts increases over time, as progenitors proliferate. Enhancing proliferation of progenitors significantly decreases oocyst numbers, while limiting proliferation increases oocyst survival. Live imaging revealed that enteroblasts interact directly with oocysts and eliminate them. Midgut progenitors sense the presence of *Plasmodium* oocysts and mount a cellular defense response that involves extensive proliferation and tissue remodeling, followed by oocysts lysis and phagocytosis of parasite remnants by enteroblasts.

**One-Sentence Summary:** Mosquito midgut stem cells detect *Plasmodium* parasites, proliferate and enteroblasts actively eliminate oocysts.

## Main text

Mosquitoes become infected with *Plasmodium* parasites when they ingest blood from a malaria-infected host. Fertilization of female gametes takes place in the gut lumen, giving rise to a motile ookinete that must traverse the midgut and can be targeted by the mosquito complement-like defense response (*1, 2*). Those ookinetes that reach the midgut basal lamina transform into oocysts, a stage in which the parasite multiplies and grows continuously for 2-3 weeks. Oocysts form a capsule and are no longer eliminated by the complement-like system (*3*). Mature oocysts rupture, releasing thousands of sporozoites that migrate to and invade the salivary gland, and are transmitted when the mosquito bites a new host. Here, we show that midgut stem cells respond to the presence of developing oocysts and proliferate to surround them and actively eliminate them. This novel stem cell-mediated defense response is modulated by the Jak/STAT pathway and results in dramatic changes in midgut architecture.

An *Anopheles stephensi* SDA-500 transgenic line (HP10) that expresses a fluorescent reporter (td-Tomato) in a subset of midgut cells with morphology reminiscent of *Drosophila* midgut stem cells (Fig. 1A) and in hemocytes (Fig.S1), was selected during an enhancer-trap screen. The *Gal4* insertion was mapped to chromosome 3 (position 22,556,999 bp, Fig S2A-C), and a single chromosomal insertion in Chr 3 (div. 41B) was confirmed by FISH hybridization (Fig S2B). The reporter protein is expressed in small triangular midgut cells in the basal plane (potential intestinal stem cells, ISCs) (Fig. 1A, front and lateral view) and in a subset of larger cells embedded in the epithelium (potential committed enteroblasts) (Fig. 1B, front and lateral view) that lack microvilli and often do not reach the surface of the gut lumen. In *Drosophila*, ISCs divide asymmetrically to produce an ISC and an enteroblasts that exits the cell cycle and differentiate into enterocytes or enteroendocrine cells (*4*). To define the identity of td-Tomato-expressing cells, mosquito midguts were stained for delta, a classic marker of pluripotency in ISC (*5*). The small triangular basal cells were positive for delta while cells embedded in the epithelia did not express *delta*, reflecting commitment to enteroblast differentiation (Fig. 1C and 1D). As expected, oral administration of bleomycin, a chemical that causes tissue damage (*6*), triggered an increase in td-Tomato positive cells (Fig. S3A and B), indicative of proliferation of midgut progenitors to restore damaged epithelial cells. Furthermore, a fraction of dissociated midgut cells enriched for td-Tomato-positive cells (Fig. S3C), with td-Tomato mRNA levels 15-fold higher than whole midguts, was also enriched for *delta* and *klumpfuss* expression, markers of midgut stem cells and enteroblasts, respectively (Fig. S3C). In contrast, expression of the *A. stephensi* ortholog of the *Drosophila* POU box gene (*Pdm-1)*, a marker of mature enterocytes (*7*), was not enriched (Fig S3C). The typical morphology, the response to chemical damage and the enrichment of gene markers, all indicate that the td-Tomato-positive midgut cells in this transgenic line are epithelial progenitor cells (stem cells and enteroblasts).

**Fig. 1:**
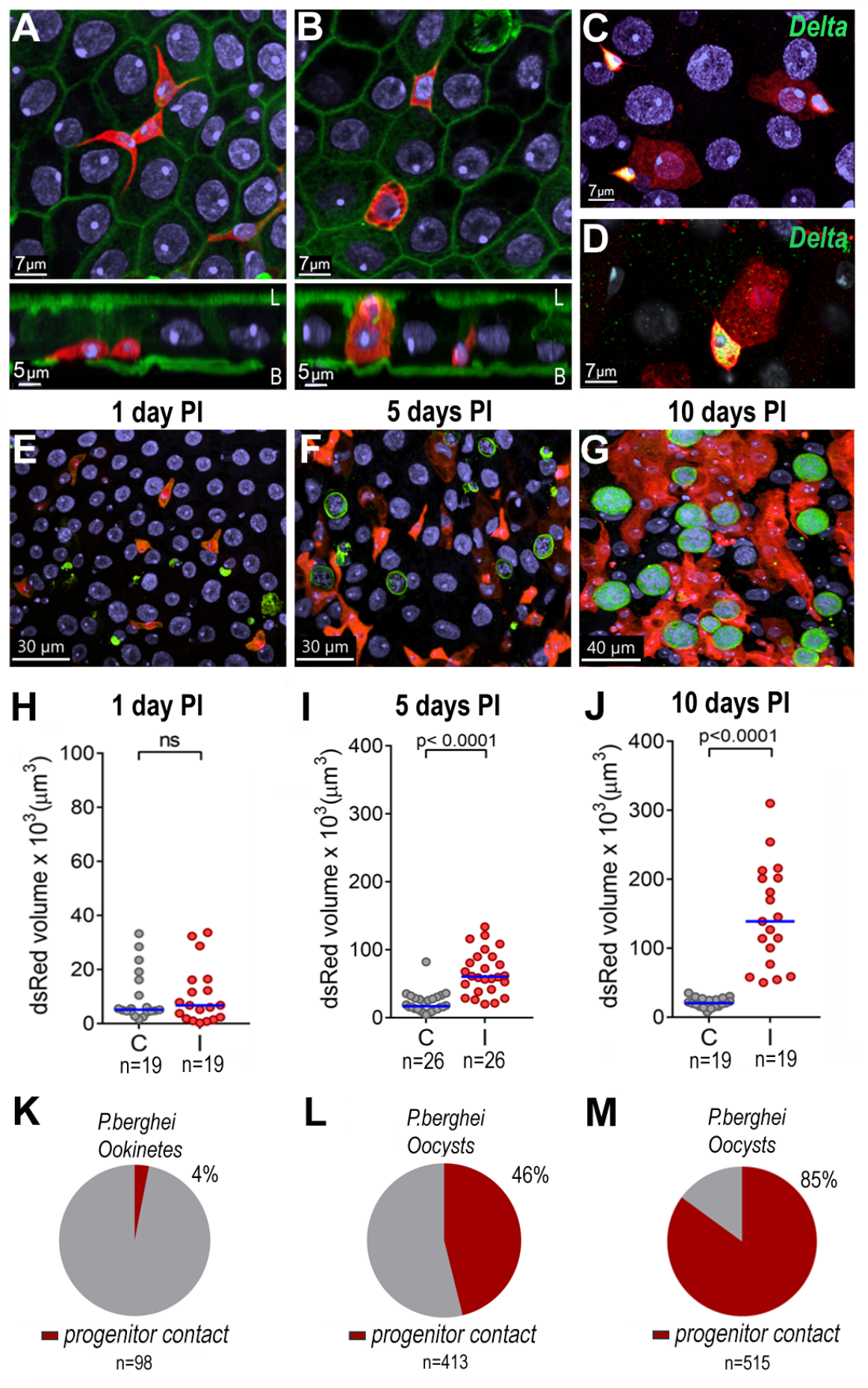
Response of *Anopheles stephensi* HP10 line midgut progenitors to *Plasmodium berghei* infection. (A) Small triangular td-Tomato^+^ cell in the basal side of the midgut epithelium. (B) Larger td-Tomato^+^ cells embedded in the epithelium. Front and lateral views. Scale bar: 7 and 5 μm, respectively. td-Tomato (red), nuclei (blue) and actin (green). L= lumen and B= basal. (C) *delta* antibody staining of small triangular td-Tomato^+^ cells. (D) Close-up of Delta immunostaining in small triangular cells. Td tomato in red; Nuclei in blue and *delta* in green. Scale bars: 7μm. Td-Tomato^+^ cells in midguts (E) 24 hours, (F) 5 days and (G) 10 days post infection (PI). Parasites (green), Td tomato (red) and nuclei (blue). Volume of dsRed (Td tomato^+^ cells) in the mosquito midgut at (H) 24 hours, (I) 5 days and (J) 10 days PI. Each dot represents the volume of red fluorescence for individual midguts and the medians are indicated with the horizontal line. Mann Whitney U test, ns = p>0.05. Percentage of *Plasmodium* parasites that are in contact with midgut progenitors at (K) 24 hours, (L) 5 days and (M) 10 days PI. C = uninfected midguts, I = infected midguts. n = number of parasites

Ookinete traversal causes irreversible damage to invaded mosquito midgut cells, which undergo apoptosis and are extruded into the midgut lumen (*8*). *P. berghei* infection resulted in no statistically significant difference in the number of midgut progenitors one day post-infection (PI) compared to uninfected midguts from blood feed females (Fig. 1E and H, Fig S4A), a time when ookinetes are traversing the midgut, and very few ookinetes (4%) were associated with progenitor cells (Fig. 1K). At 5 days PI there was a 3.5-fold increase in the number of progenitors in infected midguts (Fig. 1F and I, Fig. S4B; Mann-Whitney, p<0.0001) and 46% of oocysts were in contact with at least one progenitor (Fig. 1L). Intense proliferation and differentiation of midgut progenitors occurred between days 5 and 10 PI (Fig. 1G and J), resulting in a 6.74-fold increase in the number of progenitors in infected midguts compared to blood-fed uninfected midguts (Fig. 1J and Fig. S4C and S5; Mann-Whitney, p<0.0001). At 10 days PI most *Plasmodium* oocysts (85%) were in direct contact with midgut progenitors (Fig. 1G and M). The *Plasmodium-*induced increase in the number of enteroblasts and the displacement of epithelial cells by growing oocysts result in a dramatic reorganization of the midgut epithelium (Fig. 2 A and B). Progenitor cells proliferate forming “ribbons” of enteroblasts intercalated between mature epithelial cells (Fig. 2 B and C) that often surround and come in direct contact with developing oocysts (Fig. 2 D and F).

**Fig. 2:**
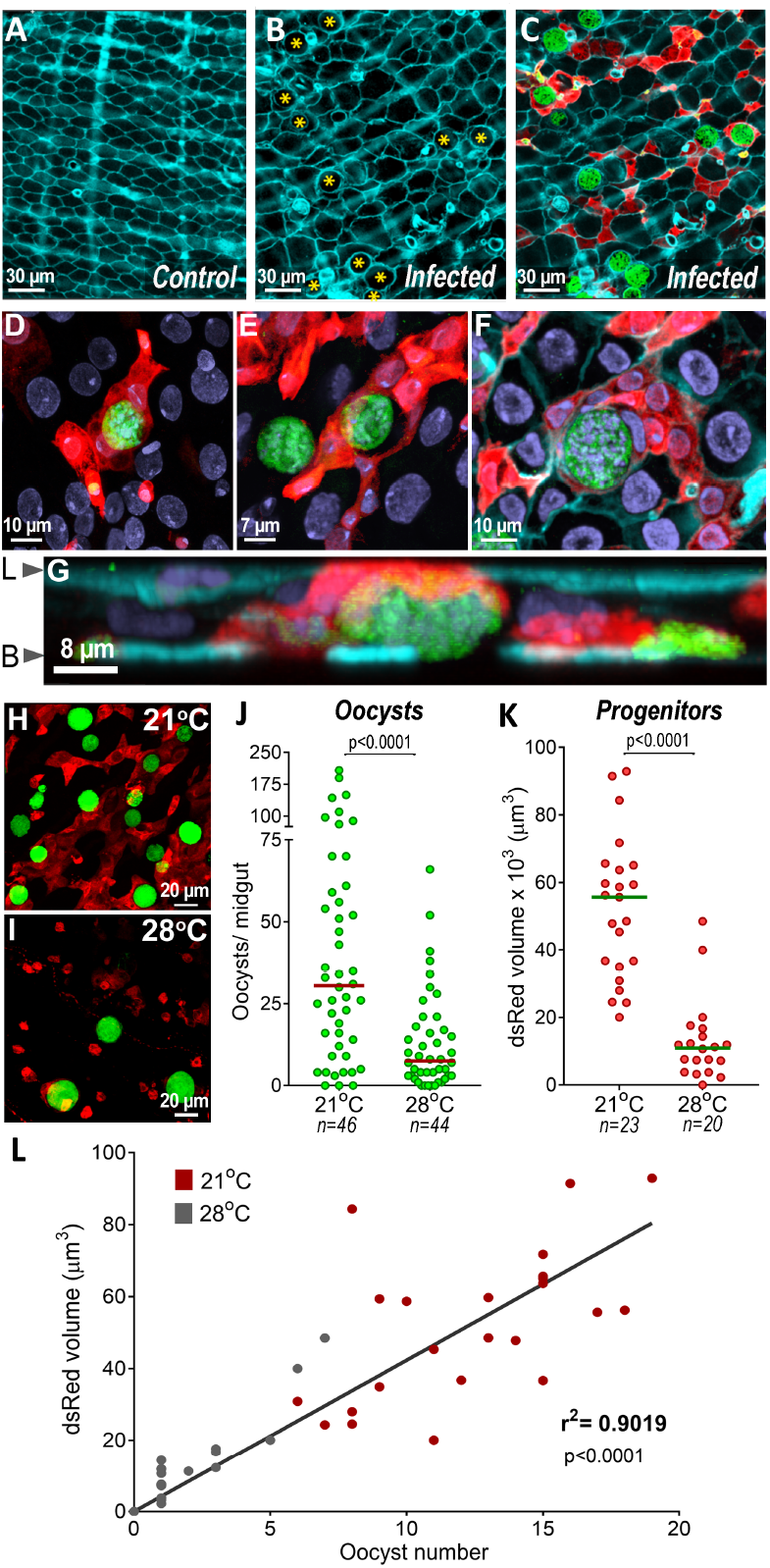
Effect of *Plasmodium* infection on midgut architecture and of oocysts intensity of infection on proliferation of midgut progenitors. (A) Actin staining of blood-fed control midguts. (B) Actin staining in *P. berghei*-infected midguts 10 days post infection. Oocysts in the epithelia are indicated by yellow asterisks. (C) Actin staining (cyan), progenitors (red) and *P.berghei* oocysts (green). Scale bar: 30μm. (D) and (E) Progenitor “ribbons” (red) surrounding oocysts (green), 10 days PI. (F) Actin (cyan) staining showing close association of progenitors (red) embedded in the epithelium with oocysts (green), 10 days PI. Scale bar: 10μm. (G) Lateral view of (F), showing midgut progenitors (red) surrounding an oocyst 10 days PI. L= lumen and B= Basal. (H) Midgut progenitors and oocysts in the midgut of *An. stephensi* kept at (H) 21°C and (I) 28°C. (J) Oocysts counts of mosquitoes kept at 21°C, a permissive temperature for oocyst development, and at 28°C, a non-permissive one. Each dot represents the number of oocysts on individual midguts. The median is indicated by the horizontal line. Mann-Whitney U test. (K) Volume of dsRed (td-Tomato^+^ cells) 10 days PI of females kept at 21°C and 28°C. Each dot represents the volume of red fluorescence for individual midguts and the medians are indicated by the horizontal line. Mann Whitney U test. (L) Correlation of oocyst numbers and dsRed volume (td-Tomato^+^ cells) of midguts from females kept at 21°C (red dots) and 28°C (gray dots). Each dot represents an individual mosquito. Linear regression, r^2^ = 0.9019.

We explored whether cell damaged by ookinete invasion was sufficient to elicit proliferation of midgut progenitors, or whether the response required the presence of developing oocysts. Two groups of mosquitoes were fed on the same infected mouse and kept at a permissive temperature (21°C) for *P. berghei* for 48h, to allow ookinete development and midgut invasion. One group was then shifted to a non-permissive temperature (28°C) 2 days PI to disrupt further oocyst development, while oocysts were allowed to mature at 21°C in the second group. As expected, the number of oocysts 10 days PI decrease significantly at a higher temperature (Mann-Whitney, p< 0.0001, Fig. 2 H-J). Although similar ookinete invasion took place in both groups, decreasing oocyst numbers significantly reduced proliferation of midgut progenitors (Fig. 2K). Furthermore, there was a strong correlation (r^2^=0.902, p<0.0001) between the volume of midgut progenitors on individual midguts and the number of oocysts, even at low oocyst density (1-20 oocysts/midgut) (Fig. 2L), suggesting that the observed proliferation of midgut progenitors is a response to the presence of oocysts.

The thioester-containing protein 1 (TEP1) is a key effector of complement-mediated elimination of *P. berghei* ookinetes in *An. gambiae* (*1*). However, silencing *LRIM1*, a gene encoding a protein that stabilizes TEP1, in *An. stephensi* SDA-500 does not significantly affect *P. berghei* survival (*9*). We confirmed that silencing *TEP1* in the *An. stephensi* HP10 transgenic line also had no statistically significant effect on parasite survival (p=0.5802, Fig. S6A), indicating that TEP1-mediated mosquito complement immunity is not limiting *P. berghei* survival. However, we documented a statistically significant decrease between the number of oocysts present at 2 and 8 days PI (Mann-Whitney, p<0.0001, Fig. S6B), suggesting that a substantial number of oocysts may be actively eliminated by mosquito defenses, as previously observed in *An. gambiae* (*10*).

Activation of JAK-STAT signaling in ISCs promotes rapid division and differentiation of progenitors (*11*). Overactivation of JAK/STAT signaling in *An. stephensi* HP10 females by silencing *SOCS* (Suppressor of Cytokine Signaling), an inhibitor of the JAK-STAT pathway, resulted in a statistically significantly increase in the number of midgut progenitors 10 days post infection (Mann-Whitney, p<0.0013, Fig. 3A). A concomitant reduction in the number of oocysts (Mann-Whitney, p<0.0004, Fig. 3B) relative to dsLacZ controls was observed, similar to what has been reported in *An. gambiae* (*10*). Conversely, disrupting JAK/STAT signaling by silencing the JAK kinase *Hopscotch* (*HOP*) greatly reduced proliferation of midgut progenitors (Mann-Whitney, p<0.0001, Fig. 3C) and significantly increased the number of oocysts (Mann-Whitney, p<0.009, Fig. 3D). Furthermore, reducing *Delta* expression promoted proliferation and differentiation of midgut progenitors (Mann-Whitney, p<0.0004, Fig. 3E), and resulted in a statistically significant decrease in oocyst numbers (Mann-Whitney, p<0.0003, Fig. 3F). Altogether, we conclude from these data that proliferation of midgut progenitors is detrimental to oocyst survival.

**Fig. 3:**
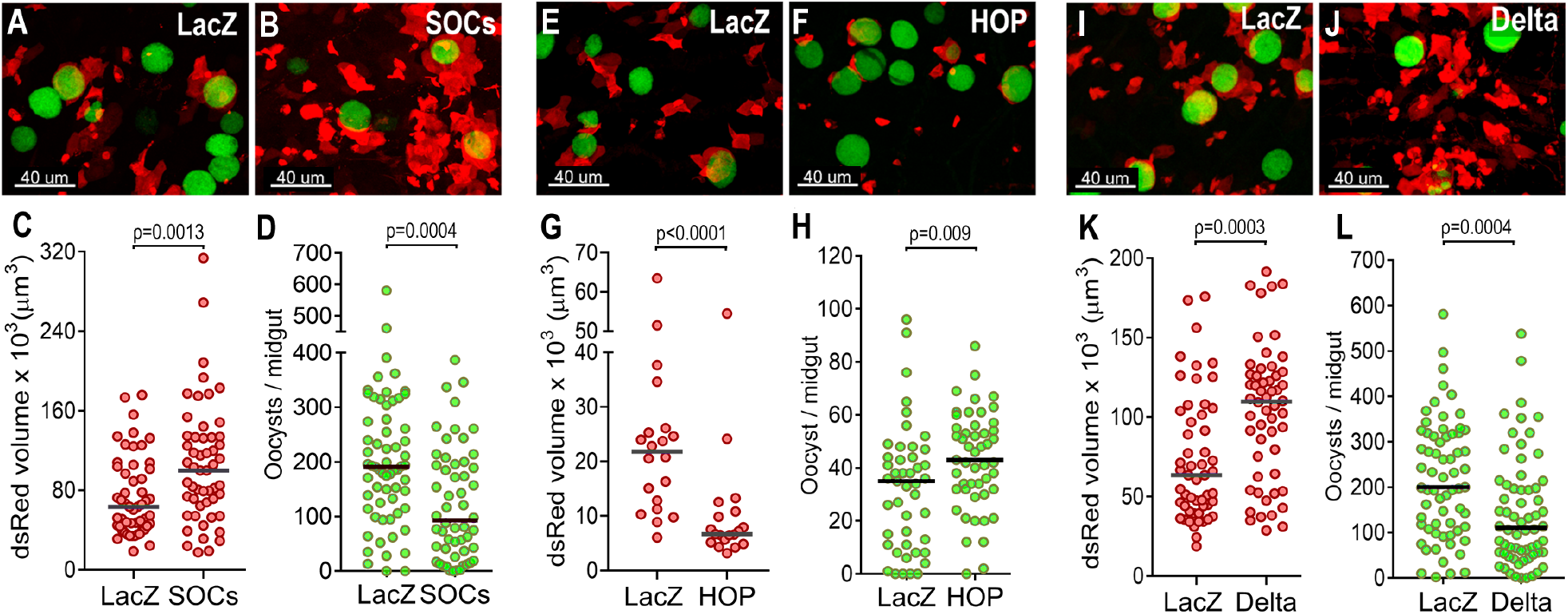
Effect of proliferation of midgut progenitors on oocysts survival. (A) *LacZ* and (B) *SOCS*-silenced midguts 10 days PI. (C) Volume of dsRed (Td tomato+ cells) in *LacZ* and *SOCS*-silenced midguts 10 days PI. (D) Oocyst counts of *LacZ* and *SOCS* silenced midguts 10 days PI. (E) LacZ and (F) HOP-silenced midguts 10 days PI. (G) Volume of dsRed (Td tomato+ cells) in *LacZ* and *HOP*-silenced midguts 10 days PI. (H) Oocyst counts of *LacZ* and *HOP-*silenced midguts 10 days po PI. (I) *LacZ* and (J) *Delta*-silenced midguts 10 days PI. (K) Volume of dsRed (Td tomato+ cells) in *LacZ* and *Delta*-silenced midguts 10 days PI. (L) Oocyst counts of LacZ and *Delta*-silenced midguts 10 days PI. Each dot represents an individual mosquito. Median is indicated by the horizontal line. Mann Whitney U test. Midgut progenitors are shown in red and *P. berghei* oocysts in green. Scale bars: 40 μm.

The hypothesis that midgut progenitors interact directly with *Plasmodium* oocysts to eliminate them was explored. Live confocal imaging was used to directly image midgut progenitor cells of live infected mosquitoes 10 and 14 days PI for periods of 9-12 hours. In females imaged 10 days PI, midgut progenitors were observed extending pseudopodia-like extensions that came in direct contact with the parasite’s surface and exerted pressure on the oocyst surface until the cell integrity was compromised and the GFP label in the oocyst cytoplasm was released (Figure 4A and Videos 1 and 2). At 14 days PI, we observed a larger oocyst being pressed on either side by two progenitor cells that also formed pseudopodia-like extensions, deforming the parasite and releasing the fluorescent cell cytoplasm towards the gut lumen (Figure 4B and Videos 3 and 4). Dead oocysts and fragments positive for GPF in immunofluorescence staining were often observed 5- and 10-days PI inside td-Tomato^+^ cells embedded in the epithelium (Fig. 4C-E and Fig S7A-C), showing that midgut enteroblasts internalize dead parasites. Furthermore, large live oocysts exhibit surface staining with the LIVE/DEAD^®^ dye, while dead oocysts are smaller, have strong signal inside the capsule and are also frequently found within midgut enteroblasts (Fig. 4F-H). At day 5 PI, 69% of oocysts in contact with midgut progenitors are dead (Table S1).

**Fig. 4:**
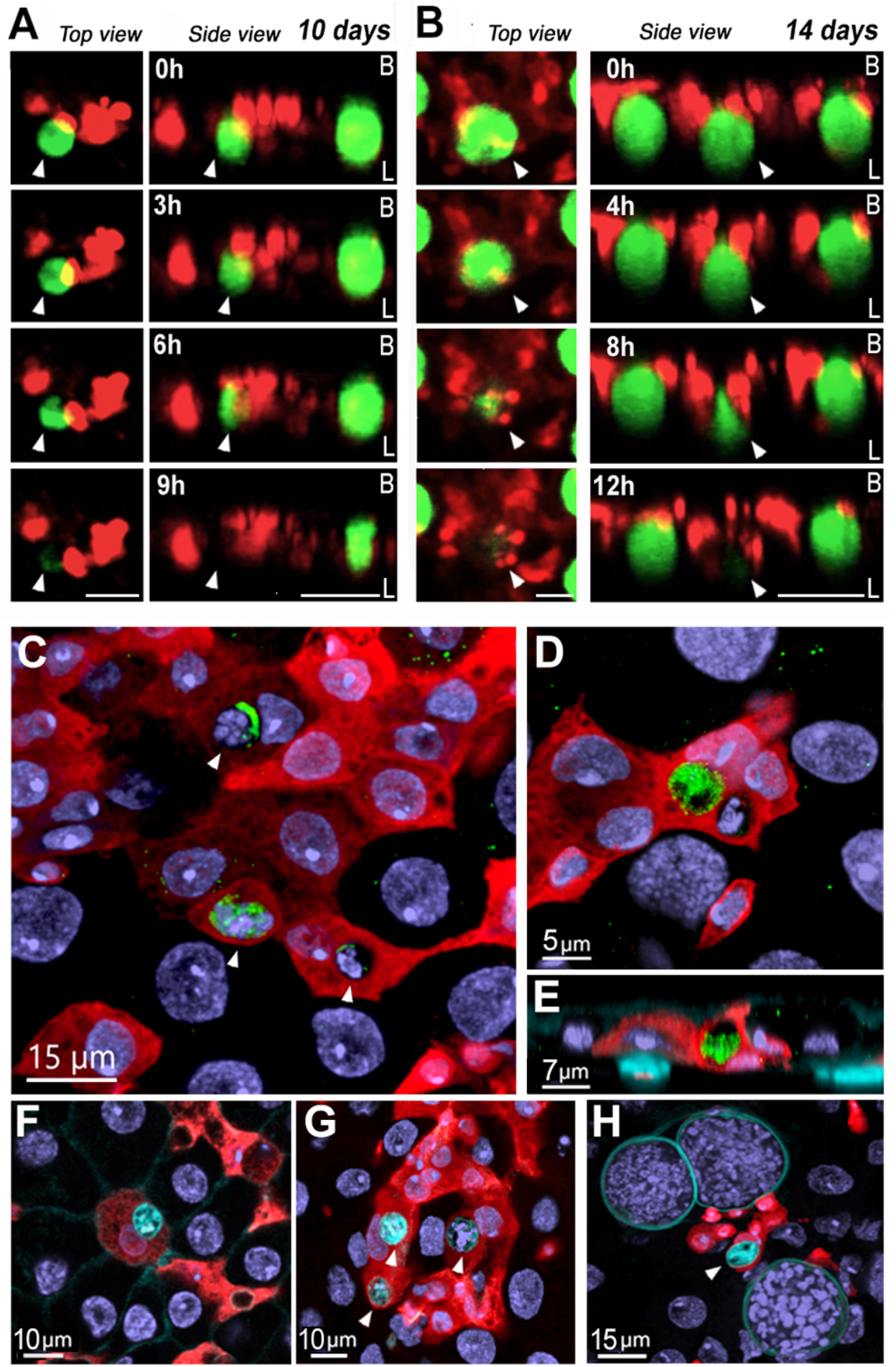
Midgut progenitors actively eliminates *Plasmodium* oocysts and internalize parasite remnants. (A) Whole mosquito live imaging over the course of 9 hours 10 days pi. Scale bar from top view: 20 μm and from side view: 30 μm. (B) Whole mosquito live imaging over the course of 12 hours 14 days PI. Scale bar from top view: 20 μm and from side view: 50 μm. (C) Oocyst fragments inside midgut progenitors at 5 days PI. White arrows indicate parasite remnants inside of midgut progenitors. Scale Bar: 15μm. Leftover of *P.berghei* GFP from inside of a midgut progenitor 5 days PI (D) Front view and (E) Lateral view. Scale Bar: 5 μm and 7 μm respectively. Midgut progenitors are in red and *P. berghei* are in green (F) Dead oocyst in cyan inside of a midgut progenitor 5 days PI. Scale Bar: 10 μm. White arrows indicate dead oocysts. (G) Dead oocysts in cyan inside midgut progenitors 5 days PI. Scale Bar: 10 μm.

*Plasmodium* infection triggered intense proliferation of midgut progenitors, which allowed enteroblasts to come in direct contact with developing oocysts. Promoting proliferation of progenitor cells by over activating STAT signaling or by reducing *Delta* expression, reduced oocyst survival. We have previously described a late-phase response mediated by the JAK/STAT pathway in *An. gambiae* with an unknown effector mechanism (*10*). Here, we provide direct evidence that, besides their critical role in maintaining the integrity of the midgut epithelial barrier in response to cell damage, stem cell-derived enteroblasts *An. stephensi* are effectors of a JAK/STAT-mediated cellular defense response against *Plasmodium* oocysts.

It is not clear how midgut progenitors detect the presence of oocysts, as initially (24h PF) they do not colocalize. One can envision that oocysts may secrete glycosyl-phosphatidyl-inositol (GPI) that are detected directly by progenitor cells. Alternatively, GPIs may activate enterocytes near the parasite and they, in turn, may release a secondary signal(s) detected by midgut progenitors. As oocysts increase in size, they may also exert physical pressure on adjacent epithelial cells and trigger the release of chemokines that promote proliferation of midgut progenitors. Live imaging indicates that midgut progenitors can move, albeit slowly, and cooperate to compress oocysts and extend pseudopod-like projections that come in direct contact with the oocysts surface before their integrity is compromised, suggesting that they release a local factor(s) that damages the oocyst capsule. Oocysts fragments are often found inside midgut enteroblasts, indicating that they internalize dead parasites. At early stages of *Drosophila* embryonic development, dying cells are mostly engulfed by sister cells that are not fully differentiated (*12*), indicating that epithelial cells with some degree of pluripotency can act as phagocytes. Taken together, our studies provide direct evidence that midgut progenitors can detect the presence of *Plasmodium* oocysts and mount a cellular defense response that involves extensive proliferation and tissue remodeling, followed by oocyst lysis and phagocytosis of parasite remnants.

## Acknowledgments

We thank Kevin Lee, Yonas Gebremicale and André Laughinghouse for insectary support.

## Funding

This work was supported by the Intramural Research Program of the Division of Intramural Research Z01AI000947, NIAID, National Institutes of Health and by NIH Grant P01GM095467.

## Author contributions

Conceptualization: ABFB, IS, FC, DO, CB-M

Methodology: ABFB, FC, SB

Investigation: ABFB, JS, EB, SB, FC

Visualization: ABFB, SB

Funding acquisition: CB-M

Project administration: CB-M

Supervision: CB-M, IS, DO

Writing – original draft: ABFB, CB-M

Writing – review & editing: ABFB, CB-M

## Competing interests

The authors declare no competing financial interests.

## Data and materials availability

All data are available in the main text or the supplementary materials.

## Supplementary Materials

Material and Methods

Figs. S1 to S7

Table S1

Movies S1-S4

## Notes

### Competing Interest Statement

The authors have declared no competing interest.

## References

1. S. Blandin et al., Complement-like protein TEP1 is a determinant of vectorial capacity in the malaria vector Anopheles gambiae. Cell 116, 661–670 (2004).

2. J. C. Castillo, A. B. B. Ferreira, N. Trisnadi, C. Barillas-Mury, Activation of mosquito complement antiplasmodial response requires cellular immunity. Sci Immunol 2, (2017).

3. R. C. Smith, C. Barillas-Mury, M. Jacobs-Lorena, Hemocyte differentiation mediates the mosquito late-phase immune response against Plasmodium in Anopheles gambiae. Proc Natl Acad Sci U S A 112, E3412–3420 (2015).

4. S. C. Herrera, E. A. Bach, JAK/STAT signaling in stem cells and regeneration: from Drosophila to vertebrates. Development 146, (2019).

5. N. Buchon, N. A. Broderick, T. Kuraishi, B. Lemaitre, Drosophila EGFR pathway coordinates stem cell proliferation and gut remodeling following infection. BMC Biol 8, 152 (2010).

6. A. Tian et al., Damage-induced regeneration of the intestinal stem cell pool through enteroblast mitosis in the Drosophila midgut. EMBO J 41, e110834 (2022).

7. J. Korzelius et al., Escargot maintains stemness and suppresses differentiation in Drosophila intestinal stem cells. EMBO J 33, 2967–2982 (2014).

8. Y. S. Han, J. Thompson, F. C. Kafatos, C. Barillas-Mury, Molecular interactions between Anopheles stephensi midgut cells and Plasmodium berghei: the time bomb theory of ookinete invasion of mosquitoes. EMBO J 19, 6030–6040 (2000).

9. P. F. Billingsley et al., Transient knockdown of Anopheles stephensi LRIM1 using RNAi increases Plasmodium falciparum sporozoite salivary gland infections. Malar J 20, 284 (2021).

10. L. Gupta et al., The STAT pathway mediates late-phase immunity against Plasmodium in the mosquito Anopheles gambiae. Cell Host Microbe 5, 498–507 (2009).

11. H. Jiang et al., Cytokine/Jak/Stat signaling mediates regeneration and homeostasis in the Drosophila midgut. Cell 137, 1343–1355 (2009).

12. Z. Zhou, P. M. Mangahas, X. Yu, The genetics of hiding the corpse: engulfment and degradation of apoptotic cells in C. elegans and D. melanogaster. Curr Top Dev Biol 63, 91–143 (2004).

13. D. A. O’Brochta, K. L. Pilitt, R. A. Harrell, 2nd, C. Aluvihare, R. T. Alford, Gal4-based enhancer-trapping in the malaria mosquito Anopheles stephensi. G3 (Bethesda) 2, 1305–1315 (2012).

14. A. B. F. Barletta, N. Trisnadi, J. L. Ramirez, C. Barillas-Mury, Mosquito Midgut Prostaglandin Release Establishes Systemic Immune Priming. iScience 19, 54–62 (2019).

15. C. J. Potter, L. Luo, Splinkerette PCR for mapping transposable elements in Drosophila. PLoS One 5, e10168 (2010).

16. M. V. Sharakhova, G. N. Artemov, V. A. Timoshevskiy, I. V. Sharakhov, Physical Genome Mapping Using Fluorescence In Situ Hybridization with Mosquito Chromosomes. Methods Mol Biol 1858, 177–194 (2019).

17. R. M. Waterhouse et al., Evolutionary superscaffolding and chromosome anchoring to improve Anopheles genome assemblies. BMC Biol 18, 1 (2020).

18. A. Amcheslavsky, J. Jiang, Y. T. Ip, Tissue damage-induced intestinal stem cell division in Drosophila. Cell Stem Cell 4, 49–61 (2009).

19. K. J. Livak, T. D. Schmittgen, Analysis of relative gene expression data using real-time quantitative PCR and the 2(-Delta Delta C(T)) Method. Methods 25, 402–408 (2001).

20. M. W. Pfaffl, A new mathematical model for relative quantification in real-time RT-PCR. Nucleic Acids Res 29, e45 (2001).

21. A. Molina-Cruz et al., Some strains of Plasmodium falciparum, a human malaria parasite, evade the complement-like system of Anopheles gambiae mosquitoes. Proc Natl Acad Sci U S A 109, E1957–1962 (2012).

22. N. Trisnadi, C. Barillas-Mury, Live In Vivo Imaging of Plasmodium Invasion of the Mosquito Midgut. mSphere 5, (2020).

