## Supplementary File for "Mosquito midgut stem cell cellular defense response limits *Plasmodium* parasite infection"

**The PDF file includes:**

Materials and Methods
Figs. S1 to S7
Table S1
Captions Movies S1 to S4
Movies S1 to S4

### Materials and Methods

Ethics statement

Public Health Service Animal Welfare Assurance #A4149-01 guidelines were followed according to the National Institutes of Health Animal (NIH) Office of Animal Care and Use (OACU). These studies were done according to the NIH animal study protocol (ASP) approved by the NIH Animal Care and User Committee (ACUC), with approval ID ASP-LMVR5.

*Anopheles stephensi HP10 hindgut transgenic line*

HP10 is a transgenic line of *Anopheles stephensi* recovered during an enhancer trap screen performed as described by O'Brochta et al. (2012) during which a *piggyBac*-based promoterless Gal4 enhancer trap element was remobilized with *piggyBac* transposase "in trans", resulting in the random remobilization of the *Gal4* containing element. Remobilized enhancer trap lines that displayed GAL4 expression specifically in adult hemocytes were established. HP10 is maintained as homozygotes at 28ºC, 80% humidity under a 12h light/ dark cycle and kept with 10% Karo syrup solution during adult stages. Tissue specific expression of *Gal4* can be visualized by indirect immunofluorescence using anti-GAL4 antibodies or by crossing HP10 to lines containing reporter genes under the regulatory control of GAL4 responsive promoters (*13*).

Mouse feeding and *Plasmodium berghei* infection

Mosquito infections with *Plasmodium berghei* were performed using transgenic *An.stephensi* HP10 mosquitoes and a transgenic GFP *P.berghei* parasite strain (ANKA GFPcon 259cl2) maintained by serial passages into 3 to 4 week old female BALB/c mice (Charles River, Wilmington, MA, USA) from frozen stocks. Infectivity was measured by parasitemia levels. Four to five-day old females were fed when mice reached 3-5% parasitemia for regular infections or before it reached 1% parasitemia for lower ones (oocyst counting experiments). Same age uninfected mice were used to feed blood-fed control mosquitoes. After feeding, both control and infected mosquitoes were maintained at 19ºC, 80% humidity and 12h light/dark cycle until the day of dissection.

Midgut Immunostaining

To image the midgut at late stages of infection, we fed mosquitos a saline solution supplemented with 10% BSA (Bovine Serum Albumin) right before dissection to distend the midgut epithelia. A day before the dissection, a cup with water was placed inside the cage for egg laying. The next day, mosquitoes were artificially fed with 10% BSA in 0.15M Sodium Chloride mixed with 10mM Sodium Bicarbonate, pH 7.2 (*14*). Sodium Bicarbonate solution must be fresh, and pH adjusted at the day of the feeding. Mosquitoes were dissected between 30 minutes and 1 hour after feeding. For one day PI timepoint midguts were already distended and mosquitoes were not fed with saline solution. After feeding, midguts were dissected in PBS at room temperature and fixed for 30 seconds with 4% Paraformaldehyde (PFA) to preserve the midgut structure. Tissues were then opened longitudinally in ice cold PBS and cleaned to remove the bolus. Then, tissues were placed in 4% PFA for at least one hour at room temperature. Tissues were then washed with 0.1% Triton PBS three times, 10 minutes each, at room temperature. Midguts were blocked for 4 hours in 2% BSA, 0.1% gelatin and 0.1% Triton in PBS at room temperature and stained overnight using either rabbit polyclonal anti-DsRed (1:1000) (Living Colors® DsRed Polyclonal Antibody, 632496, Takara Bio, San Jose, CA, USA) or mouse monoclonal anti-RFP (1:1000) (Rocklands Immunochemicals, Philadelphia, PA, USA) to visualize cells that expressed d-Tomato depending on antibody combination. To visualize ookinetes (1 day PI) in the midgut we stained midguts using mouse anti-Pbs21 and to stain early oocysts (5 days PI) we used rabbit polyclonal anti-PbCap380

(1:1000). To visualize oocysts at later stages (10 days PI) we used its inner GFP fluorescence and to detect oocysts fragments inside the enteroblasts we incubated tissues with rabbit anti-GFP (1:500) (Anti-GFP antibody, ab6556, Abcam, Cambridge, UK). For Delta staining, midguts were incubated with mouse monoclonal anti-Delta (1:20) (Dl antibody, C594.9B, DSHB, Iowa City, IA, USA) in blocking solution (PBS containing 2% BSA, 0.1% gelatin and 0.1% triton). All primary antibody incubations were performed overnight at $4^0$C.

After primary antibody, midguts were washed with PBS containing 2% BSA, 0.1% gelatin and 0.1% triton and incubated with secondary antibodies goat anti-rabbit (1:1000) (Alexa Fluor 488 or 594, Molecular Probes, ThermoFisher Scientific, Waltham, MA, USA) or goat anti-mouse (1:1000) (Alexa Fluor 488 or 594, Molecular Probes, ThermoFisher Scientific, Waltham, MA, USA) for 2 hours at room temperature. Midguts were washed three times with blocking buffer and washed twice with PBS with 0.1% triton. Tissues were incubated with 20μM Hoechst 33342 (405, Molecular Probes, ThermoFisher Scientific, Waltham, MA, USA) for nuclei staining and 1U of phalloidin (Alexa Fluor 647 or Alexa Fluor 750, Molecular Probes, ThermoFisher Scientific, Waltham, MA, USA) to visualize the midgut actin, diluted in PBS with 0.1% triton for 30 minutes at room temperature. Microscope slides were mounted using a drop of Prolong Gold Antifade Mountant (Molecular Probes, ThermoFisher Scientific, Waltham, MA, USA).

dsRed volume quantification and surface analysis

To determine the volume of the cells expressing td-Tomato within the midgut we used the surface mode on Imaris 9.9.1 (Bitplane, Concord, MA, USA). To create a surface, we used confocal z-stack sections. First, we applied the gaussian filter to smooth the picture, then we used the surface mode with a threshold of 16 for the number of voxels. After the surfaces were generated, we applied a refining filter to remove smaller volumes and eventual tracheal signal. From the surface created from the td-Tomato signal we were able to quantify the volume of the midgut progenitors in the tissue. To determine the association between midgut progenitors and the parasite at different stages of infection. We also created a surface for the parasite fluorescent signal. We smooth the picture as described before and then created a surface with a threshold of 45 for number of voxels. Small surfaces were eliminated with a second filter to keep only the surfaces correspondent to parasites. We counted the number of parasite surfaces and how many were associated with the surfaces of midgut progenitors.

<u>Midgut oocyst counting</u>

*Plasmodium berghei* infections were evaluated by counting oocyst numbers per mosquito midgut after feeding on an infected mouse. Infected mosquitoes were kept at 19ºC for 10 days after feeding when they were dissected, and their midgut fixed in 4% PFA for 15 minutes at room temperature. After washing with PBS three times, midguts were mounted in a slide and counted under a fluorescence microscope, where live oocysts were identified by their GFP expression. The number of oocysts per midgut was represented in a scatter plot where each dot represents an individual mosquito. For the experiment to count oocysts 2 and 8 days after infection (Fig S5B), we used a *P.berghei* mCherry strain which is brighter than GFP and easier to count at early days without the need of antibody staining. At 2 days post infection, fed mosquitoes kept at 19ºC still have a blood bolus, therefore they were opened and cleaned before fixation with 4% PFA, as described above. For oocysts counting, midguts were mounted using VECTASHIELD® Antifade Mounting Medium with DAPI (H-1200-10, Vector Laboratories, Newark, CA, USA).

<u>Confocal microscopy</u>

Confocal images were captured using a Leica TCS SP8 (DM8000) confocal microscope (Leica Microsystems, Wetzlar, Germany) with either a 40x or a 63x oil immersion objective equipped with a photomultiplier tube/ hybrid detector. Midguts were visualized with a white light laser, using 498-nm excitation for Alexa 488 (phalloidin or *P.berghei* parasites); 588-nm excitation for Alexa 594 (midgut progenitors); 644-nm excitation for Alexa 647 (Live/Dead probe and phalloidin) and a 405-nm diode laser for nuclei staining (Hoechst 33342). Images were taken using sequential mode and variable z-steps. Image processing and merge was performed using Imaris 9.9.1 (Bitplane, Concord, MA, USA) and Adobe Photoshop CC (Adobe Systems, San Jose, CA, USA).

Mapping *Gal4* enhancer-trap insertion

Splinkerette PCR was used to map the genomic location of the *Gal4* enhancer-trap insertion. Briefly, genomic DNA was isolated from individual *An. stephensi* HP10 larvae (4$^{th}$ instar). The genomic DNA was digested by BstYI to produce sticky ends. A double stranded splinkerette oligonucleotide with stable hairpin loop and compatible sticky ends is ligated to the digested genomic DNA (SPLNK-GATC-TOP GATCCCACTAGTGTCGACACCAGTCTCTAATTTTTTTTTCAAAAAAA and SPLNK-BOT – CGAAGAGTAACCGTTGCTAGGAGAGACCGTGGCTGAATGAGACTGGTGTCGACACT AGTGG). Next, we perform two rounds of nested PCR. For the first round we used the following primers: SPLNK#1- CGAAGAGTAACCGTTGCTAGGAGAGACC and 3’SPLNK-PB#1 – GTTTGTTGAATTTATTATTAGTATGTAAG (to map 3 prime end) or 5’SPLNK-PB#1- ACCGCATTGACAAGCACG (to map 5 prime end). For the second round we used the following primers: SPLNK#2 – GTGGCTGAATGAGACTGGTGTCGAC and 3’SPLNK#2 – GGATGTCTCTTGCCGAC (to map 3 prime end) or 5’SPLNK-PB#2 – CTCCAAGCGGCGACTGAG (to map 5 prime end). Those two rounds generate a PCR fragment that contains the flanking genomic DNA between the *piggybac* element insertion site and the genomic digestion site. Detailed protocol is described in (*15*). PCR fragments generated from the 3 and the 5 prime ends were cloned in TOPO-TA vector (K450002, ThermoFisher Scientific, Waltham, MA, USA) and then used for a standard Sanger sequencing reaction using TOPO vector primers.

Fluorescence *in situ* hybridization of the GAL4-specific fluorescent probe with the polytene chromosomes of *An. stephensi* HP10 transgenic line

Fluorescence in situ hybridization (FISH) was done following the previously published protocol with minor modifications (*16*).

Fluorescent DNA probe preparation

Genomic DNA was extracted from *An. stephensi* HP10 hindgut mosquitoes and amplified with *Gal4*-specific primers (Gal4clone_F – AAGAAAAACCGAAGTGCGCC and Gal4clone_R – CACCAAACAAAGCAGACGGG). About 30 ng of the purified PCR product were used in a Random-primer labeling reaction, in which Cyanine 5-dUTP (Enzo Life Sciences Inc., Ann Arbor, MI, USA) was incorporated into the DNA by Klenow fragment (ThermoScientific, Graiciuno, Lithuania) and Random Primers DNA Labeling System (Invitrogen, Carlsbad, CA, USA). The manufacture-supplied protocol for the Random Primers DNA Labeling System was used. The labeled DNA was precipitated by 2.5 volumes of 96% Ethanol and 0.1 volume of 3M sodium acetate at −20°C overnight and then centrifuged at 14,000 rpm, +4 °C for 20 min. The supernatant was removed and the pellet of DNA was air-dried and dissolved in 50 µl of hybridization buffer (60% deionized formamide, 2× SSC, 10% dextran sulfate) by shaking the mix in the Eppendorf ThermoMixer C (MilliporeSigma, St. Louis, MO, USA) at 2,000 rpm, 37 °C for 1 hour.

Polytene chromosome preparation

*An. stephensi* HP10 females were fed with blood twice after emergence and allowed to lay eggs. A third blood meal was offered, and females were allowed to develop their ovaries for 26 hours at 26°C and 80% humidity. Ovaries were dissected from half-gravid females under a MZ6 Leica stereo microscope (Leica Camera, Wetzlar, Germany) and fixed in fresh modified Carnoy's solution: methanol:glacial acetic acid (3:1). The ovaries were kept at room temperature overnight until they were transferred ovaries to -20 °C for a long-term storage. Parts of dissected ovaries were put in drops of 50% propionic acid on slides for about 5 minutes until follicles become clear and about twice of their original size. Follicles were separated from each other with needles, and other tissues were removed by wiping them away with a piece of a paper towel. A fresh drop of 50% propionic acid was applied to the separated follicles. Follicles were covered with a dust-free coverslip and left for about 5 minutes. A piece of filter paper was placed over the coverslip that was gently tapped with a pencil eraser to release polytene chromosomes from the nurse cells of follicles. The banding pattern and spreading of polytene chromosomes was examined using an Olympus CX43 Phase Microscope (Olympus, Tokyo, Japan) with a 20× objective. Slides with suitable chromosomal preparations were placed in a humid chamber with 4× SSC in the bottom of the chamber and incubated at +4 °C overnight for better flattening of chromosomes. Slides were immersed in liquid nitrogen until the bubbling stops (10–15 seconds). After taking the slides out of the liquid nitrogen, the coverslips were removed with a razor blade. Slides were immediately placed in a slide jar with prechilled 50% ethanol (−20 °C) and kept at +4 °C for at least 2 hours. The preparations in a slide jar were dehydrated with ethanol series of 70% and 90% for 5 minutes each at +4 °C and then with 100% ethanol for 5 minutes at room temperature. Slides were air-dried and kept in a box until the hybridization.

Fluorescence *in situ* hybridization

Slides with chromosomal preparations were incubated in the 2× SSC with 4% formaldehyde solution at 60 °C for 30 min. Then, 15 µl of the labeled GAL4-specific fluorescent DNA-probe dissolved in the hybridization buffer were applied to the slide and covered with a 22×22 coverslip. The borders of the coverslip were insulated with rubber cement. To denature DNA of chromosomes and the probe, the slides were then put into the Thermobrite machine (Leica Biosystems, Wetzlar, Germany) and heated at +80 °C for 5 min. Hybridization occurred at +37 °C overnight. On the next day, the slides were washed in 2× SSC at 60°C for 15 min, then in 2× SSC at room temperature for 15 min, and finally in 0.2× SSC for 10 min. The washing solution was removed from the slide and 15 µl of Prolong Gold Antifade Mounting with DAPI (ThermoFisher Scientific, Waltham, MA, USA) was applied to the slide and covered with a 22×22 coverslip. Microscopy and image acquisition were performed with an Axio Imager Z1 microscope (Carl Zeiss MicroImaging GmbH, Munich, Germany). Image processing was performed by the Fiji software (cite: doi:10.1038/nmeth.2019). Mapping of the location of the probe was done according to the standard cytogenetic map of polytene chromosomes for *An. stephensi* (*17*).

Midgut chemical damage by Bleomycin feeding

To induce chemical damage in the midgut, 3- to – 4 day old females of *An.stephensi* HP10 hindgut line were fed with 10% Karo syrup solution supplemented with 25 µg/ml Bleomycin (Bleomycin sulfate, anticancer and antibiotic agent, ab142977, Abcam, Cambridge, UK) for 2 days (*18*). Fresh solution was offered every day of the treatment. After 2 days, females were artificially fed with 10% BSA saline solution to distend the midgut tissue as described above.

Statistical analysis

All statistical analyses were performed using GraphPad Prism 5 (GraphPad, San Diego, CA, USA). All analyses where pertinent were conducted using either the student Unpaired t-test, Mann-Whitney Test, ANOVA Dunnett multiple comparison and ANOVA Kruskal-Wallis test and are indicated in the legend of each figure when appropriate. The test choice was determined based on the variation of the sample, if parametric or non-parametric, and the number of treatments in each experiment. Significance was assessed at $p < 0.05$. The error bars represent the Standard Error of the Mean (SEM).

Enrichment of midgut progenitors for RNA extraction

*An. stephensi* HP10 hindgut females were dissected, and 15 midguts were transferred to a tube containing 100 µl of elastase solution in PBS (1 mg/ml) (07453, Stem Cell Technologies, Vancouver, British Columbia, Canada) to dissociate the midgut tissue. Incubate the tubes in a shaker for 1 hour at 27°C, pausing to pipette up-and-down every 15 minutes with a siliconized pipette tip to avoid tissues to stick. After incubation, 100 µl of 2% BSA solution in PBS was added to the mix to stop digestion. Digested tissue was then loaded into a 20-µm cell strainer (pluriStrainer Mini 20 µm, pluriSelect, Leipzig, Germany). We passed the filtrate two more times on the same filter with a new pipette tip every time. We added 900 µl of TRIzol LS (TRIzol™ LS

Reagent, 10296010, ThermoFisher Scientific, Waltham, MA, USA) and proceeded to RNA extraction. We compared the RNA expression of midgut enriched fractions with RNA from whole midguts placed straight into 1ml of TRIzol.

RNA extraction, cDNA synthesis and qPCR analysis

Pools of 15 whole midguts from *An.stephensi* HP10 line were collected from sugar-fed females and placed directly into 1ml TRIzol reagent (TRIzol™ reagent, 15596026, ThermoFisher Scientific, Waltham, MA, USA) and homogenized with a motorized pestle. The midgut enriched fractions were placed in TRIzol LS as described above and RNA extraction was conducted as follows. Two hundred microliters of chloroform were added (1/5 of TRIzol volume) to 1mL of TRIzol and vortex vigorously for the aqueous phase separation. Samples were centrifuged for 12,000 RCF, 10 minutes, 4$^{o}$C. The aqueous phase was then transferred to a fresh 1.5 mL Eppendorf tube and the RNA precipitated by adding 0.25 mL of isopropyl alcohol (500 mL per 1 mL TRIZOL reagent used). Five microliters of linear acrylamide (5 mg / mL) were also added to each tube to aid in precipitation and pelleting. Samples were mixed by repeated inversion 10 times, incubated for 10 minutes at room temperature, and then spun at 12,000 RCF, 10 minutes, 4$^{o}$C. All the supernatant was removed, and the RNA pellets washed twice with 75% ethanol (minimum 1 mL of ethanol per 1 mL of TRIZOL used). To wash the pellets, tubes were mixed by vortexing and centrifuged 7,500 RCF, 5 minutes, 4$^{o}$C. After the last wash supernatant was removed and samples air-dried until almost dry, but not completely (still translucent). RNA was solubilized with 30 µL of RNAse-free water, pipetting a few times to homogenize and then placed at 55$^{o}$C for 10 minutes to resuspend the RNA. RNA concentration was measured at 260nm using a Denovix Spectrophotometer (DS-11 Series Spectrophotometer / Fluorometer). One microgram of total RNA was used for complementary DNA (cDNA) synthesis using Quantitect Reverse Transcription

Kit (Qiagen, Germantown, MD, USA) according to manufacturer's instructions. Gene expression was assessed by quantitative PCR (qPCR) using the resulting cDNA as a template. Quantitative PCR (qPCR) was used to measure td-Tomato (AY678269.1), *delta* (ASTE009642 or ASTEI20_036824), *klumpfuss* (Klu) (ASTE009884 or ASTEI20_041957) and *POU domain transcription factor* (Pdm) (ASTE011391 or ASTEI20_034725) gene expression in whole sugar-fed midguts and fractions enriched for midgut progenitors. We used the DyNamo SYBR green qPCR kit (ThermoFisher Scientific, Waltham, MA, USA) with specific primers and the assay ran on a CFX96 Real-Time PCR Detection System (Bio-Rad, Hercules, CA, USA). A 133-bp fragment was amplified for td-Tomato (F- ATCGTGGAACAGTACGAGCG and R- TGAACTCTTTGATGACGGCCA). A 165-bp fragment was amplified for *delta* (F- TGGGAGTTTCAACCGACTGG and R- CGATCGGTGAGCAGGTGTAA). A 154-bp fragment was amplified for *Klu* (F- AGTCTCCACAGCAACCGATG and R- CGGGCAAACTCCTGGTAGAG). A 134-bp fragment was amplified for *Pdm* (F- GCCTATCCTCACCTTCGTCC and R- CGGTCATTCCTGCTTGATGC). Relative expression was normalized against *An. stephensi* ribosomal protein S7 (RpS7) as internal standard and analyzed using the ΔΔ Ct method (*19, 20*). RpS7 (ASTE004816 or ASTEI20_033711) primers sequences were: F- TGGAAATGAACTCGGATCTGAAG and R – CCTTCTTGTTGTTGAACTCGACCT. Statistical analysis of the fold change was performed using Unpaired t-test (GraphPad, San Diego, CA, USA). Each independent experiment was performed with three biological replicates (three pools of 15 mosquitoes) for each condition.

dsRNA synthesis and injection for gene silencing

Three-to-four day old female *An.stephensi* HP10 females were cold-anesthetized and injected with 69nl of a 3µg/µl *dsSOCS, dsHOP, dsDelta* or *dsLacZ* control. Double-stranded RNA for

*Suppressors Of Cytokine Signaling* (SOCS) (ASTE009458 or ASTEI20_041097), *Hopscotch* (HOP) (ASTE008682 or ASTEI20_034867) and *delta* (ASTE009642 or ASTEI20_036824) was synthesized by *in vitro* transcription using the MEGAscript RNAi kit (Ambion, ThermoFisher Scientific, Waltham, MA, USA). DNA templates were obtained by PCR using *An.stephensi* cDNA extracted from whole body sugar-fed females. A 431-bp fragment was amplified for SOCS with primers containing T7 promoters (F-**TAATACGACTCACTATAGGG** TTCATCCACTGTCTGGTGCC and R- **TAATACGACTCACTATAGGG** TTTGGTAGCGTCAGCTCGTT), using an annealing temperature of 58$^{o}$C. A 498-bp fragment was amplified for HOP with primers containing T7 promoters (F-**TAATACGACTCACTATAGGG**CGATGGTGCTAGAATTTCCG and R-**TAATACGACTCACTATAGGG**CGCAGCTCAAACACTCGTAG), using an annealing temperature of 58$^{o}$C. A 305-bp fragment was amplified for *delta* with primers containing T7 promoters (F-**TAATACGACTCACTATAGGG**GCGGTCAGTCTTGTGAGGAA and R-**TAATACGACTCACTATAGGG**TTCGTTCTGCTTTCTCGCCT), using an annealing temperature of 58$^{o}$C. Double-stranded RNA for *LacZ* was synthetized by amplifying a 218-bp fragment from *LacZ* gene clones into pCRII-TOPO vector using M13 primers to generate a dsRNA control as previously described (*21*). Injected mosquitoes were fed in a *P.berghei*-infected mouse to evaluate the effect of gene silencing in the oocyst number. Silencing efficiency with this method ranged between 50%-60% for all the genes tested in the midgut.

*In vivo* live imaging

*An.stephensi* HP10 females infected with *P.berghei* were allowed to lay eggs and then fed another blood meal to distend the midgut. After the second meal mosquitoes were placed at 19$^{o}$C again. Mosquito abdomens must be distended to allow visualization of the midgut through the cuticle (transparent lateral panel). We used mosquitoes at two timepoints during infection, 10 and 14 days post *P.berghei* infection. Imaging took place the next day at 18-20 hours post-bloodmeal. Mosquitoes were imaged as previously described (*22*). In short, we removed legs and head of 5-10 mosquitoes and were placed between a coverslip and glass slide with craft putty as a spacer. Images were taken on a Leica SP8 confocal microscope using a 40x 1.25 NA oil objective with a white light laser with excitation at 561 nm for td-Tomato and at 488 nm for GFP-expressing *P.berghei* oocysts. The z-stack with 1 µm intervals was taken every 10 minute for 9-12 hours with resonant scanner. Videos were processed using Imaris 9.9.1 (Bitplane, Concord, MA, USA) and saved as Tiff series. Then files were imported to ImageJ and then converted to the MOV format.

<u>Viability of oocysts in the midgut</u>

To access the viability of the oocysts in the midgut, we artificially fed a 10% BSA saline solution described above to engorge the midgut and preserve the morphology for microscopy. Shortly after feeding *P.berghei*-infected midguts from *An. stephensi* HP10 females were dissected and quickly fixed for 30 seconds in 4% PFA at room temperature, they were then placed in a well containing ice-cold PBS. When a pool of 10 midguts was reached, tissues were transferred to well containing a LIVE/DEAD™ Fixable Far Red Dead Cell Stain (L34973, ThermoFisher Scientific, Waltham, MA, USA) solution diluted 1:1000 in PBS. Midguts were incubated in the Live/Dead stain solution for 30 minutes on ice. After that, the tissues were opened longitudinally to form a single sheet, as described above. The bolus content was removed and washed out and tissues were placed in 4% PFA fixative solution at room temperature for at least an hour. Finally, midguts were submitted to an immunofluorescence protocol to add antibodies to stain either midgut progenitors, described above. Live oocysts presented a weaker fluorescent signal in their capsule on the outside, while dead oocysts had a brighter signal inside. Live/Dead fluorescent stain was evaluated on a Leica

SP8 confocal microscope using either a 40x or a 63x oil objective depending on the size of the oocyst with a white light laser with excitation at 633-635 nm for the Live/Dead fluorescent stain. Live/Dead signal is preserved even with the use of detergents present during the immunofluorescence protocol.

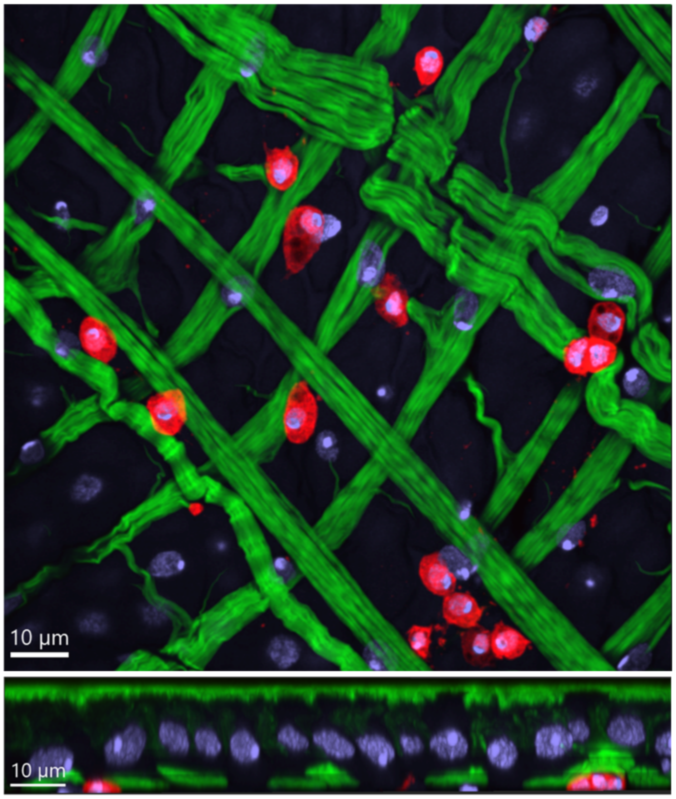


**Fig. S1. An *Anopheles stephensi* SDA-500 transgenic line (HP10) that expresses a fluorescent reporter (td-Tomato) also in hemocytes.** XY and lateral views of hemocytes attached to the midgut surface. Hemocytes are located on top of the muscle layer showed in green. Hemocytes in red, nuclei in blue and actin in green. Scale bar: 10 µm.

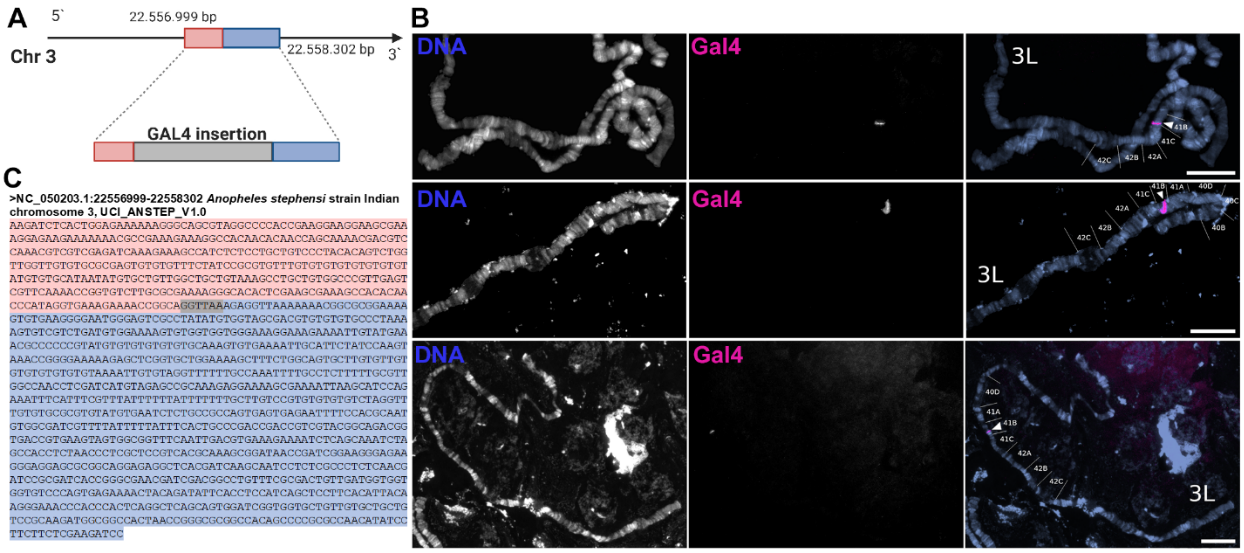


**Fig. S2. A single GAL4 enhancer-trap insertion was mapped to chromosome 3.** (A) Schematic drawing representing the insertion site of the *Gal4* enhancer trap in the HP10 transgenic line. (B) FISH to detect GAL4 with ovarian nurse cell polytene chromosomes. A single *Gal4* insertion is shown by arrows on chromosome 3 (3L arm; subdivision 41B). First column indicated DNA staining; middle column shows the *Gal4* insertion, and the last column is a merged image. Scale bar: 20 µm. (C) Region of *piggyBac* insertion in the genome. The sequences flanking the enhancer mapped to chromosome 3 in the region between 22.556.999 bp and 22.558.302 bp is shown. The region 5' prime from the insertion is highlighted in pink and the 3' prime flanking region in blue. The six nucleotides highlighted in gray are present on both sides of the insertion and therefore represent the potential insertion site of the enhancer.

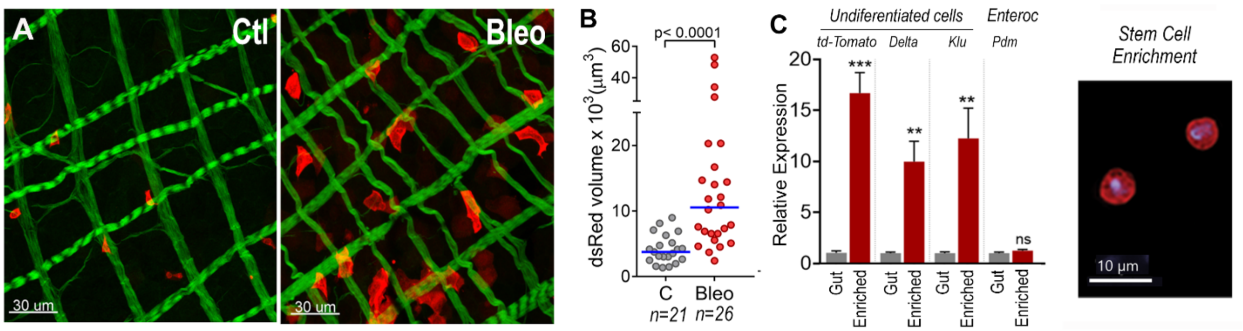


**Fig. S3. Td-tomato$^+$ midgut cells from *An.stephensi* HP10 transgenic line are epithelial progenitors (stem cells and enteroblasts).** (A) Midgut surface of mosquitoes treated with control sugar solution or 25µg/ml bleomycin solution for 48 hours. Actin muscle layer is in green, and progenitors are in red. Scale bar: 30 µm. (B) Volume of dsRed (td-Tomato$^+$ cells) in control sugar-fed and bleomycin-treated midguts, 2 days post-treatment. Each dot represents an individual midgut. Median is shown as a blue horizontal bar. Mann Whitney U test. (C) Gene expression of td-Tomato, *Delta*, *Klumpfuss* and *Pdm* in whole sugar-fed midguts and fractions enriched for midgut progenitors. Error bars represent mean ± SEM of midgut pools. Unpaired t-test, **P≤0.01, ***P≤0.001.

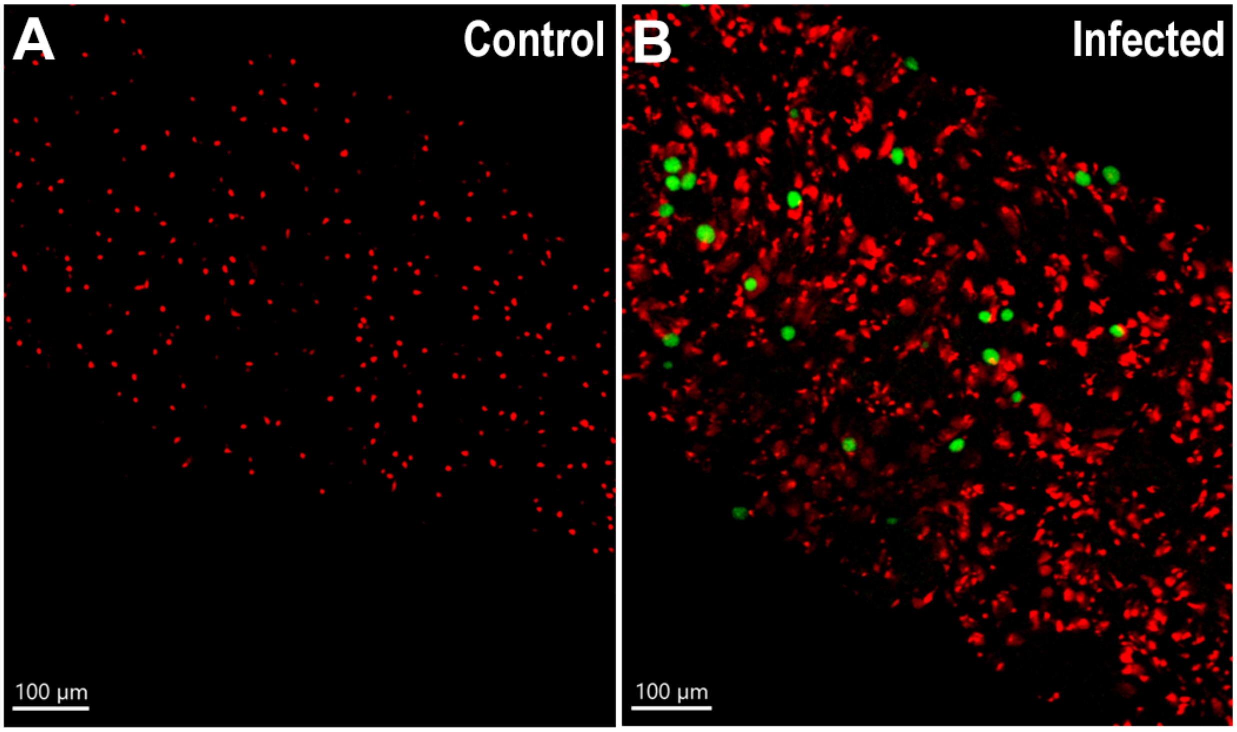


**Fig. S4. Midgut progenitors proliferate more in infected midguts and form clusters around oocysts.** (A) Midgut surface highlighting midgut progenitors in blood fed mosquitoes, 10 days post feeding (one side of the midgut layer); 10.5 µm section. (B) Midgut surface highlighting midgut progenitors in *P. berghei* infected mosquitoes 10 days post feeding (one side of the midgut layer); 10.5 µm section. Midgut progenitors are shown in red and *Plasmodium* oocysts in green. Scale bar: 100 µm.

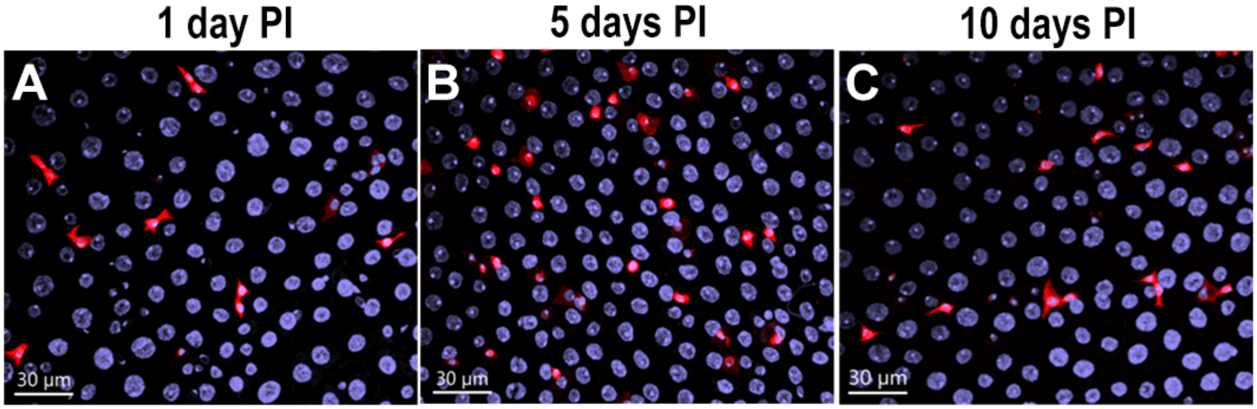


**Fig. S5. Midgut progenitors in blood fed mosquitoes at different time points after feeding.** (A) Midgut surface highlighting midgut progenitors in blood fed mosquitoes, (A) 1 day; (B) 5 days; and (C) 10 days post feeding. Midgut progenitors are shown in red and nuclei is in blue. Scale bar: 30 µm.

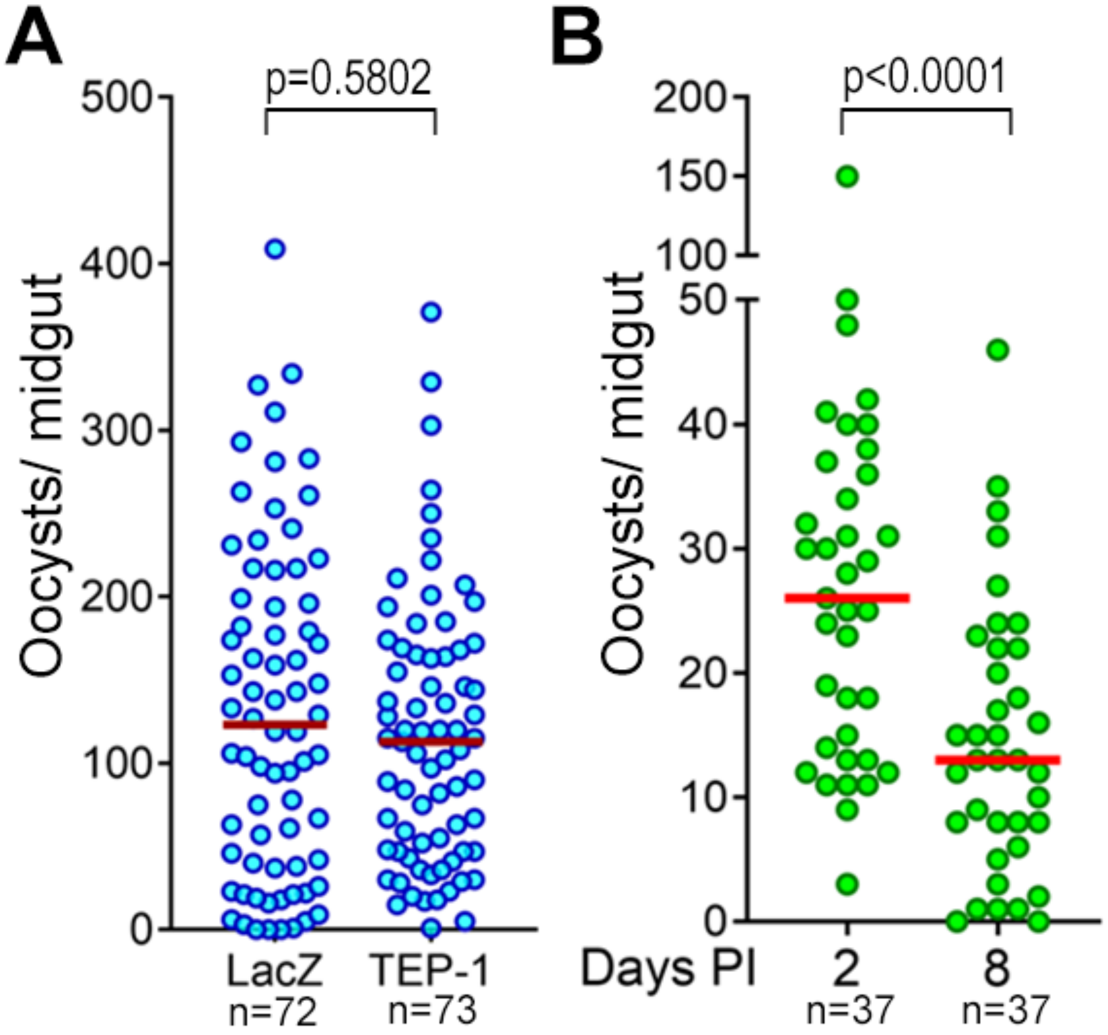


**Fig. S6. *Anopheles stephensi* eliminates oocysts using a late-phase immune mechanism.** (A) Oocysts counts of LacZ and TEP-1-silenced midguts 10 days post *P.berghei* infection. (B) Oocysts counts of *An.stephensi* infected with *P.berghei* at day 2 and day 8 post-infection. Each dot represents an individual mosquito. Median is shown as red horizontal bars. Mann Whitney U test.

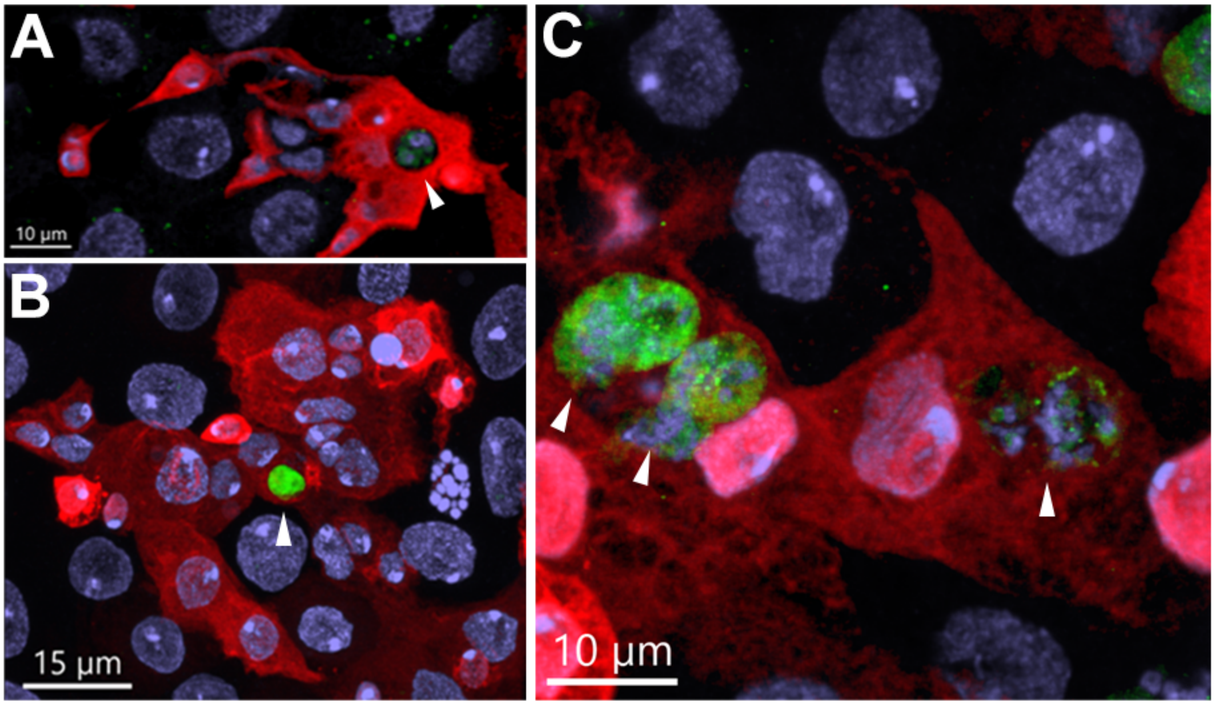


**Fig. S7. Midgut progenitors internalize oocysts fragments.** (A) Oocyst found inside of a midgut progenitor. Oocyst nuclei are still visible. Scale Bar: 10 µm. (B) GFP from oocysts can be found inside midgut progenitors. Scale bar: 15 µm. (C) Close-up of pieces of oocysts inside midgut progenitors. Nuclei and GFP signal can be seen. Scale bar: 10 µm. White arrows indicate internalized parasites. Midgut progenitors are shown in red and *P.berghei* in green.

| Groups | Exp 1 | Exp 2 | Compiled | Percentage % |
| --- | --- | --- | --- | --- |
| **Associated parasites - Live** | 19 | 11 | 30 | 31 |
| **Associated parasites - Dead** | 43 | 23 | 66 | 69 |
| **Total** | 62 | 34 | 96 | 100 |

**Table S1. Percentage of dead oocysts associated with midgut progenitors in the midgut.**

**Movie S1.**

Midgut progenitors in direct contact eliminate *P.berghei* oocysts from the midgut epithelia 10 days after infection (Top view).

**Movie S2.**

Midgut progenitors in direct contact eliminate *P.berghei* oocysts from the midgut epithelia 10 days after infection (Side view).

**Movie S3.**

Midgut progenitors in direct contact eliminate *P.berghei* oocysts from the midgut epithelia 14 days after infection (Top view).

**Movie S4.**

Midgut progenitors in direct contact eliminate *P.berghei* oocysts from the midgut epithelia 14 days after infection (Side view).
